# A Time-Resolved High-Throughput Screening of Fission Yeast Deletion Mutants for Oxidative Stress Resistance

**DOI:** 10.64898/2025.12.22.695924

**Authors:** Mohammadtaha Pirsalehi, Rowshan Ara Islam, Kristal Ng, Olga Xintarakou, Peter Thorpe, Charalampos Rallis

## Abstract

Cells typically balance growth with stress responses - growing rapidly in low stress conditions and halting growth to defend against stress or to repair stress-induced damage. While numerous genome-wide screens have identified mutants resistant to oxidative stress, these have largely relied on static, end-point measurements. Here, we take a dynamic, time-resolved approach to uncover how fission yeast, *Schizosaccharomyces pombe*, adapts to oxidative stress over time. We have tracked the growth of 3,420 deletion mutants across nine time points spanning four days on both nutrient-rich solid media and that containing oxidative stress induced by hydrogen peroxide. This kinetic strategy revealed not just resistant or sensitive mutants. It allowed clustering of growth patterns across time and uncovered mutants that are capable of transiently uncoupling growth from stress response. Hydrogen peroxide induced a dose-dependent delay in colony expansion in most deletion strains, yet 15 mutants consistently maintained robust growth. These belong to different functional categories, highlighting diverse potential mechanisms ranging from altered DNA damage checkpoints to metabolic rewiring and growth regulation. By capturing dynamic trajectories rather than static outcomes, this study exposes hidden layers of growth under oxidative stress and identifies new genetic determinants of cellular resilience in fission yeast.

## Introduction

Reactive oxygen species (ROS) are inevitable byproducts of aerobic metabolism and can also arise from environmental stressors such as UV light, ionizing radiation, and hydrogen peroxide (H_2_O_2_) [1, 2]. When present in excess, ROS overwhelm the cellular antioxidant capacity, causing oxidative stress that damages DNA, proteins, and lipids. This oxidative damage contributes to ageing and underlies many pathologies, including cancer and neurodegenerative diseases[3, 4]. In biotechnological contexts, oxidative stress also impairs yeast productivity by reducing viability, growth rate, and metabolic efficiency. Yet, ROS are not merely toxic byproducts: at physiological levels, they serve essential signalling roles in growth, metabolism, and stress adaptation. Thus, maintaining ROS homeostasis is critical for sustaining cellular function [5, 6].

The fission yeast *Schizosaccharomyces pombe* is a powerful eukaryotic model for dissecting conserved stress-response mechanisms. With a compact genome organised into only three chromosomes and more than 70% of its protein-coding genes having clear metazoan orthologues, *S. pombe* provides a tractable and evolutionarily relevant system for exploring fundamental principles of cell biology [7]. Its fully annotated genome and comprehensive gene-deletion libraries make it particularly suitable for genome-wide screening and systems-level analysis [8, 9].

Adaptation to stress requires precise reprogramming of gene expression. *S. pombe* integrates nutrient and stress signals primarily through two evolutionarily conserved pathways: the Target of Rapamycin (TOR) and Stress-Activated Protein Kinase (SAPK) cascades [10–13]. The TOR complex 1 (TORC1) promotes growth in response to amino acid and nutrient availability [14–17]. whereas SAPKs are activated under adverse conditions such as oxidative, nutritional, heat, osmotic and other stresses to promote defence mechanisms and repair. These pathways act antagonistically: when growth-promoting TORC1 activity is high, stress protection is suppressed, and vice versa. In fluctuating environments, cells must continuously recalibrate this balance to optimise proliferation while preserving resilience [2, 10, 11].

During environmental stress, *S. pombe* mounts two levels of protection: a Core Environmental Stress Response (CESR) and stress-specific responses. The CESR, also termed the global stress response, reprograms transcription to divert cellular resources from growth toward protection and recovery, and is activated by diverse stressors, including heat, osmotic, and oxidative stress [18]. In *S. pombe*, CESR activation is largely mediated by the Sty1-Atf1 mitogen-activated protein kinase cascade. Under oxidative stress, ROS indirectly activate Sty1, which phosphorylates the transcription factor Atf1, triggering the induction of genes required for repair and detoxification [1, 19–21]. In parallel, *S. pombe* employs oxidant-specific regulators such as the transcription factors Pap1 and Prr1. Pap1 mediates defence against low levels of H_2_O_2_ independently of Sty1-Atf1, but under high oxidative load it becomes coupled to the pathway via Srx1, a sulfiredoxin that reactivates oxidised Tpx1 -the direct activator of Pap1 [22–25]. Prr1 acts independently of both Sty1 and Pap1 to control a subset of oxidative stress response genes [1, 26].

Despite extensive study of oxidative stress signalling in *S. pombe*, hydrogen peroxide resistance has never been systematically examined as a primary phenotype in a genome-wide screen. Previous reports have only identified H_2_O_2_-resistant or -sensitive mutants as secondary outcomes within broader phenotypic surveys [27, 28]. Two main strategies exist for genome-wide screening in *S. pombe*: pooled barcoded mutant libraries in liquid culture and arrayed mutants on solid media [9, 29–31]. However, liquid-culture approaches can produce artefacts due to competition between strains of differing growth rates [32], while most solid-media studies have relied on endpoint imaging, missing the dynamic recovery phase following stress exposure [33]. Because many stress-induced transcripts return to near-baseline levels even under continued stress [2, 34], such endpoint assays can yield false positives or negatives.

To overcome these limitations, we conducted a dedicated, time-resolved genome-wide deletion screen for hydrogen peroxide resistance in *S. pombe* using arrayed mutants on solid agar. Colonies were imaged repeatedly over four days, allowing us to capture dynamic growth trajectories rather than static outcomes. Beyond identifying resistant strains, our aim was to discover mutants that could uncouple growth from stress response: cells that sustain or even accelerate growth under oxidative challenge.

This question is biologically and biotechnologically important. In general, cells that grow slowly exhibit longer lifespans and greater stress tolerance [35–37], reflecting the antagonism between TOR-driven growth and stress resistance [38–40]. To test whether oxidative stress-resistant mutants follow this paradigm, we performed chronological lifespan (CLS) assays, which measure the survival of non-dividing cell populations in stationary phase [41].

Identifying and characterising such resistant mutants has dual value: it provides new insight into the genetic architecture of oxidative stress defence and reveals strains with potential industrial applications. Because stress-tolerant yeasts can thrive under harsh production conditions, they represent promising platforms for biomanufacturing of biofuels, pharmaceuticals, and therapeutic proteins such as monoclonal antibodies [42–44].

## Results

### Genome-wide screening of *S. pombe* deletion mutants exposed to H_2_O_2_

To systematically identify the genetic determinants that alter resistance to oxidative stress, we performed a genome-wide screen of *Schizosaccharomyces pombe* deletion mutants[45]. A total of 3,420 non-essential gene deletion strains were arrayed on YES agar and challenged with hydrogen peroxide (H_2_O_2_). We initially evaluated a range of H_2_O_2_ concentrations (2–8 mM). However, these doses resulted in minimal growth inhibition, likely due to cooperative stress tolerance within high-density colony arrays, where communal factors can enhance oxidative stress resistance. Based on preliminary titration experiments, 10 mM and 12 mM H_2_O_2_ were selected as optimal concentrations that induced pronounced growth delay in most mutants, while still allowing differential growth among resistant strains.

The complete deletion collection was arrayed at a density of 384 colonies per plate across nine plates per condition. Plates were imaged at nine timepoints over a 96-hour period (0, 20, 24, 44, 48, 68, 72, 92, and 96 hours), and colony growth was quantified using Gitter, an established R-based image analysis pipeline for high-throughput colony phenotyping [46]. Visual inspection confirmed marked growth impairment under both oxidative stress conditions (Fig. 1A). To enable robust comparison across timepoints and strains, colony areas were normalized to initial (0 h) colony sizes, yielding growth trajectories across the time course (data available in supplemental Tables 1 and 2). Median growth of H_2_O_2_-exposed colonies diverged from untreated controls as early as 20 h and remained significantly attenuated thereafter (Fig. 1B), validating that 10 mM and 12 mM H_2_O_2_ elicited sustained oxidative stress.

**Figure 1.**
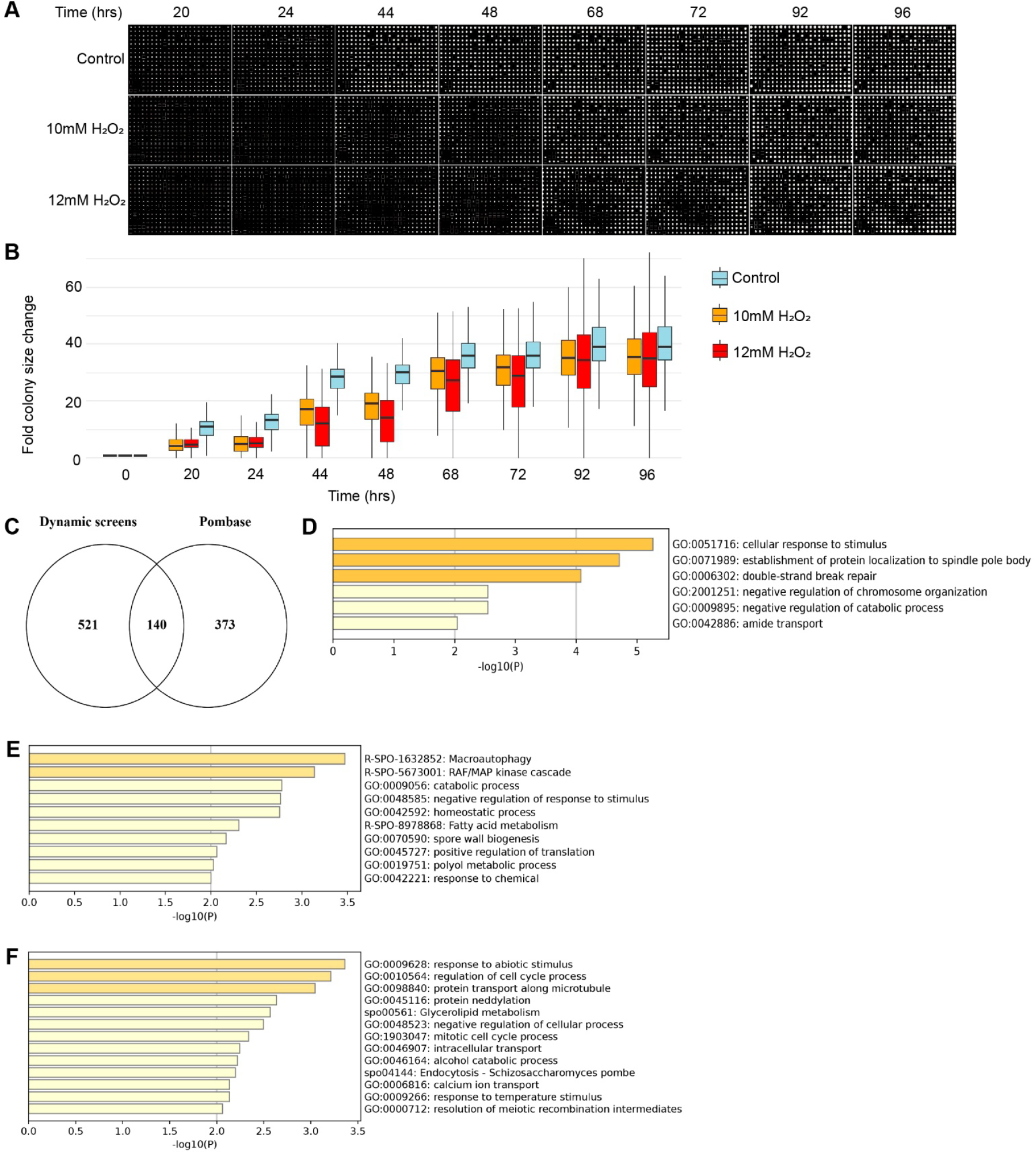
High-throughput dynamic colony growth profiling identifies oxidative stress–responsive deletion mutants. **(A)** Time-course images of colony growth for the deletion mutant library under control, 10 mM H_2_O_2_, and 12 mM H_2_O_2_. Plates were monitored across the indicated time points (20–96 h). Oxidative stress treatment resulted in delayed growth and reduced colony size compared to untreated controls, with the strongest effect at 12 mM H_2_O_2_. **(B)** Quantification of fold change in colony size across time for all deletion strains under control, 10 mM H_2_O_2_, and 12 mM H_2_O_2_ conditions. Boxplots represent the distribution of colony growth trajectories at each time point, revealing a progressive divergence in growth dynamics under oxidative stress relative to controls. **(C)** Venn diagram comparing oxidative stress-responsive candidates identified in this dynamic screen with annotated oxidative stress-associated genes in Pombase. A subset of 140 genes overlapped between datasets, indicating that the time-resolved colony growth approach captures both known and previously uncharacterized oxidative stress modulators. **(D–F)** Gene Ontology (GO) biological process enrichment analysis for strains exhibiting condition-specific growth behaviors. **(D)** GO terms enriched among mutants demonstrating enhanced growth under oxidative stress relative to control. Enriched functions include cellular response to stimulus, protein localization to the spindle pole body, and DNA repair pathways. **(E)** GO terms enriched in mutants showing altered growth dynamics in the control condition, including macroautophagy, MAPK cascade signaling, metabolic regulation, and translation-associated processes. **(F)** GO terms enriched among mutants with peroxide-sensitive growth profiles, including cell cycle regulation, intracellular transport, endocytosis, calcium ion transport, and meiotic recombination resolution. Together, these analyses highlight functional modules associated with oxidative stress adaptation and growth control.

As anticipated, the majority of mutants exhibited impaired growth relative to untreated controls, reflected by resistant ratio values <1. The resistance ratio is calculated by dividing each mutant’s colony size on the H_2_O_2_ plate by its colony size on the control plate, showing how well it grows under stress relative to normal conditions. Thus, a resistance ratio of 1 implies that a mutant is not affected by hydrogen peroxide treatment. Notably, growth inhibition correlated with the dose of H_2_O_2_: 12 mM H_2_O_2_ elicited significantly lower resistance ratios than 10 mM, as assessed by the Wilcoxon rank-sum test (p < 0.003) (Fig.1B). We defined oxidative-stress-resistant mutants as those with a stringent resistance ratio >1.10 under either H_2_O_2_ condition. Such an applied cutoff is considered stringent as it identifies mutants that are not adversely affected by the treatment and may even exhibit enhanced growth relative to control conditions. 661 deletion strains exhibited enhanced growth in the presence of oxidative stress (Supplemental Table 3). We compared this high-confidence resistance gene set to previously annotated oxidative-stress-resistant mutants in the *S. pombe* database, Pombase [47]. While there was substantial overlap, we also identified numerous strains uniquely detected by our dynamic growth-rate-based approach as well as strains previously reported but not recovered in our screen (Fig. 1C). This incomplete concordance is consistent with prior observations that phenotypic screens, even when targeting identical stress responses, often display substantial variability [48]. Gene Ontology (GO) enrichment analysis revealed distinct functional signatures across shared and unique mutant sets. Shared resistant strains were enriched in categories including spindle pole body organization, double-strand break repair, negative regulation of chromosomal architecture, catabolic pathways, and amide transport (Fig. 1D). Mutants uniquely annotated in Pombase were enriched for macroautophagy, diverse catabolic processes, and cellular homeostasis pathways (Fig. 1E). In contrast, mutants uniquely identified in our time-resolved screen showed enrichment for abiotic stress response pathways, cell-cycle regulatory proteins, and intracellular trafficking functions (Fig. 1F). Together, these analyses support the sensitivity and complementarity of real-time growth profiling in uncovering oxidative stress resilience mechanisms beyond previously catalogued phenotypes.

In summary, our results expand the oxidative stress resistome of *S. pombe*, providing dynamic phenotypic data and revealing novel genetic regulators of H_2_O_2_ tolerance. These findings highlight the modular and multifaceted nature of oxidative stress resilience and establish a resource for probing conserved stress-response pathways relevant to lifespan regulation and cellular homeostasis.

### Mutants that uncouple growth from oxidative stress response

Using longitudinal colony size measurements, we performed k-means clustering to classify deletion mutants based on their growth dynamics under control and oxidative stress conditions (12 mM H_2_O_2_). K-means clustering was conducted in the Morpheus[49], software package. This analysis yielded seven distinct groups (Fig.2A) showing marked heterogeneity in growth responses, with several clusters exhibiting highly variable and noisy trajectories. By contrast, mutants in control media displayed largely uniform growth (Fig. 2B).

**Figure 2.**
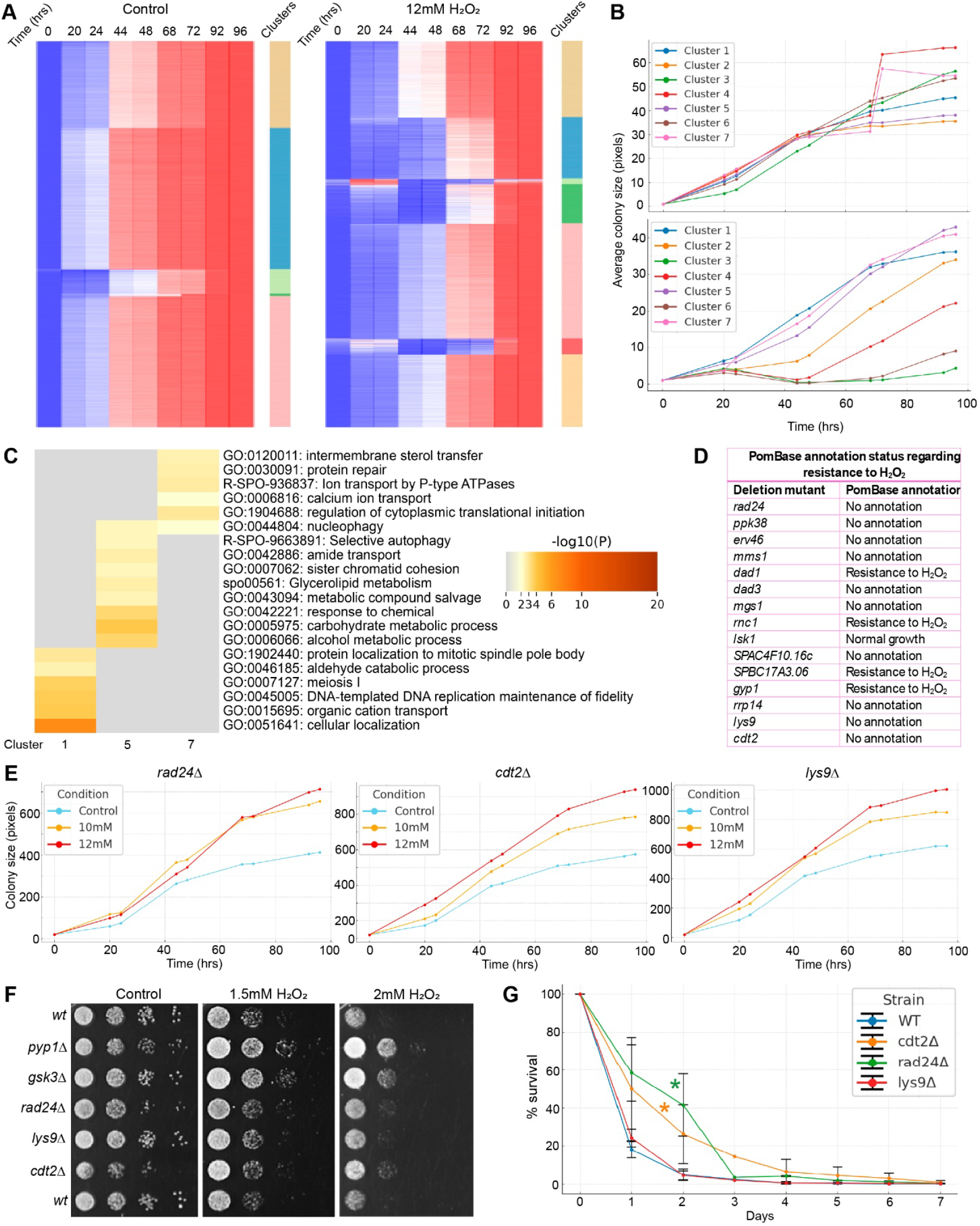
Dynamic clustering identifies oxidative stress-specific growth behaviours and mutants with enhanced stress tolerance. **(A)** Heatmap of colony growth trajectories for all deletion strains under control and 12 mM H_2_O_2_. Colours indicate fold-change in colony size (0-96 h). Seven growth clusters were identified per condition, with greater phenotypic diversity emerging under oxidative stress. **(B)** Mean growth curves of the seven clusters. Control conditions show largely uniform growth, whereas 12 mM H_2_O_2_ yields distinct and variable trajectories. Clusters 1, 5, and 7 exhibit accelerated growth under stress. **(C)** GO enrichment for fast-growing stress clusters (1, 5, 7). Enriched pathways include membrane transport, protein repair, autophagy, translation regulation, and metabolism, indicating that disruption of these functions may enhance growth under oxidative challenge. **(D)** Annotation status of 15 candidate mutants selected based on stress-enhanced growth. Only four were previously noted as H_2_O_2_-resistant in Pombase, highlighting discovery of uncharacterized stress-responsive genes. **(E)** Individual growth time-courses for *rad24Δ, cdt2Δ*, and *lys9Δ* under control, 10 mM, and 12 mM H_2_O_2_. All three mutants show enhanced proliferation compared to wild type under stress. **(F)** Spot assays validating oxidative resistance. *rad24Δ, cdt2Δ*, and *lys9Δ* display increased tolerance up to 2 mM H_2_O_2_ relative to wild type; *pyp1Δ* and *gsk3Δ* serve as positive controls. **(G)** Chronological lifespan analysis. *rad24Δ* and *cdt2Δ*, but not *lys9Δ*, exhibit modest lifespan extension relative to wild type. Stress tolerance and survival are not uniformly coupled.

We annotated each cluster according to its mean growth-curve slope, categorizing them as “slow,” “normal,” or “fast” growers. Clusters 1, 5, and 7 represented the fastest-growing groups (Fig. 2B). Gene Ontology (GO) enrichment analysis [50] revealed distinct biological themes: cluster 1 was enriched for genes involved in cellular localization, cluster 5 for carbohydrate metabolism, and cluster 7 for regulation of cytoplasmic translational initiation (Fig. 2C). These results suggest that loss of genes governing specific metabolic, trafficking, or translational processes may confer a proliferative advantage under oxidative stress. To identify stress-adaptive mutants, we focused on strains whose growth was accelerated in H_2_O_2_ (Fig. 2B). Fifteen mutants met these criteria (Fig. 2D), yet only four had been previously annotated as H_2_O_2_-resistant. Three of these strains without prior oxidative stress annotations and diverse functions, one related to DNA damage -*rad24Δ*, one to lysine metabolism *-lys9Δ*, and one to cell cycle -*cdt2Δ*, were followed up for further validations. Colony growth curves confirmed enhanced growth under peroxide, supporting the predictive value of our clustering strategy (Fig. 2E). To independently validate oxidative stress resistance, we conducted serial dilution spot assays with fast-growing cultures at OD_600_=0.5. Unlike the genome-wide screen, which required higher peroxide concentrations due to mass-protection effects on solid media, validation assays used lower concentrations, enabled by pre-growth in liquid culture. This approach allowed the use of peroxide concentrations commonly employed in the field [48]. Positive controls (*pyp1Δ* [51] and *gsk3Δ* [28]) were included. All three candidate mutants exhibited increased tolerance up to 2 mM H_2_O_2_, whereas the wild type showed limited growth at 1.5-2 mM (Fig.2F).

There are links between ageing and oxidative stress[52]. We have previously shown that, in contrast to slow growth, resistance to oxidative stress is not necessarily linked to long chronological lifespan in fission yeast[53]. We, therefore, wondered whether our fast-growing (within stress conditions) mutants exhibit altered lifespan. Colony Forming Units (CFUs) were quantified across seven days in stationary phase. Wild-type survival dropped below 1% by day 5, and *lys9Δ* showed a similar decline. In contrast, *rad24Δ* and *cdt2Δ* maintained survival above 1% until day 6, indicating a modest maximal lifespan extension. In addition, *rad24Δ* and *cdt2Δ* exhibit statistically significant (*: p<0.01, Fig. 2G) median lifespan increase. Thus, while oxidative stress resistance is not universally associated with longevity, loss of *Rad24* or *Cdt2* links DNA-damage and cell-cycle regulatory pathways to improved stress tolerance and extended lifespan. Together, these results identify a subset of deletion mutants with enhanced oxidative stress tolerance without a growth rate compromise. Our screen prioritized mutants that maintained or accelerated proliferation under peroxide challenge. Our findings underscore that distinct genetic perturbations may promote both stress resistance and a growth advantage under challenge conditions. Decoupling stress resistance and responses and growth inhibition has important implications for understanding adaptive responses to oxidative stress.

## Discussion

We used a genome-wide deletion library to identify *S. pombe* genes whose loss promotes resistance to hydrogen peroxide. Exposure to 10 mM and 12 mM H_2_O_2_ delayed the growth of most deletion mutants; only a minority maintained growth, while others failed to grow at all. We identified 15 candidate mutants resistant to hydrogen-peroxide stress and able to grow fast. Out of these 15, we further analysed three, *Δrad24, Δlys9*, and *Δcdt2*, that might potentially uncouple growth from the stress response. The validation assay demonstrated that the mutants were resistant to H_2_O_2_. In addition, GO enrichment of faster-growing clusters under 12 mM H_2_O_2_ highlighted deletions in genes involved in cellular localization, carbohydrate metabolic processes, and regulation of cytoplasmic translational initiation, suggesting that loss of these functions can facilitate faster growth under oxidative stress. In CLS assays, *rad24Δ* and *cdt2Δ* exhibited extended lifespan compared to wild type, whereas *lys9Δ* did not, indicating that oxidative stress tolerance does not necessarily correlate with altered lifespan.

Our observations align with the core environmental stress response modules (CESR) in yeasts. Acute oxidant exposure induces antioxidant and repair functions while repressing ribosome biogenesis and other growth programs, a shift that depends on dose and time and is largely regulated by the Sty1-Atf1 MAPK pathway in *S. pombe* (and *Pap1* at lower ROS). Even under persistent stress, cells adapt to stress, and growth eventually resumes, explaining the recovery phase we observed between 48–68 h on plates (ratios trending back toward 1) [1, 2, 54]. As confirmed by our validation assay, *rad24Δ, lys9Δ, and cdtΔ2* are resistant to H_2_O_2_. The “enhanced growth” seen on plates likely may reflect colony-level uncoupling of growth from stress response in these conditions. Dense communities can detoxify H_2_O_2_ and lower the effective dose for interior cells. Yeast colonies are known to behave differently in communities. This difference in H_2_O_2_ exposure between agar and liquid cultures explains why colonies can appear to outgrow stress while liquid cultures do not [55, 56].

GO analysis point to three biological processes whose deletion could improve growth under H_2_O_2_. First, deletions that shift carbohydrate metabolism away from glycolysis can divert carbon into the oxidative pentose-phosphate pathway (PPP), boosting NADPH production to fuel glutathione and peroxiredoxin systems; key glycolytic steps or oxidant-mediated inhibition of glycolytic enzymes GAPDH/PK, shift towards PPP, and improve oxidant resistance [57]. Second, under oxidative stress, cells downregulate cytoplasmic translation initiation via eIF2α phosphorylation and TORC1 inhibition, conserving energy and limiting damage. Thus, deleting positive regulators of initiation can result in a growth advantage when cells are exposed to hydrogen peroxide [58, 59]. Finally, cellular localization refers to the processes that transport a molecule, protein complex, or organelle to a defined site within the cell. We also know that *Sty1*-Atf1 localization is critical upon stress, the MAPK Sty1 accumulates in the nucleus and cooperates with Atf1p to initiate stress-gene transcription. Therefore, we hypothesize that deleting genes that facilitate cellular localization could disrupt *Sty1* trafficking; if so, the usual downregulation of TORC1 may fail to occur, consistent with reports that stress inhibits TORC1 in fission yeast and, as a result, cells grow faster [58–60].

We validated that *rad24Δ, lys9Δ*, and *cdt2Δ* are resistant to H_2_O_2_. *lys9* encodes saccharopine dehydrogenase, an enzyme involved in lysine biosynthesis; loss of *lys9* causes lysine auxotrophy. Importantly, the rich YES medium includes lysine as a supplement, so *lys9*Δ cells have access to external lysine. In budding yeast, adding extracellular L-lysine triggers “lysine harvesting”; Cells import far more lysine than they need for growth, which allows them to save NADPH and boost glutathione (GSH). This re-routing increases cellular GSH and improves resistance to oxidants. We hypothesize that *lys9Δ* causes cells to import abundant lysine from YES, saving NADPH for antioxidant systems and enhancing H_2_O_2_ tolerance [61, 62].

*rad24*, a 14-3-3 protein, binds the mitotic activator Cdc25 and helps the G2/M checkpoint; when Cdc25 is phosphorylated by checkpoint/stress kinases, *rad24* keeps it away from the nucleus to delay mitosis. Based on this checkpoint logic, we hypothesize that *rad24Δ* mutants fail to arrest in G2 during oxidative stress, rather than improving H_2_O_2_ detoxification, they keep dividing despite DNA damage [63, 64].

Cdt2 is the substrate receptor of the CRL4-Cdt2 E3 ubiquitin ligase. In *S. pombe*, CRL4-Cdt2 is recruited when PCNA is loaded on DNA during S-phase or repair and targets specific proteins for ubiquitylation and degradation. By removing these targets, CRL4-Cdt2 prevents re-replication and helps maintain adequate dNTP pools. Loss of *cdt2* leaves these proteins in place, extends S-phase, activates checkpoints, and often slows proliferation [65, 66]. In our liquid assay, *cdt2Δ* showed slow growth; cells likely spent longer in S-phase. We propose that this S-phase extension causes the H_2_O_2_ resistance, consistent with the general trend that a slow growth rate contributes to oxidative stress tolerance [37].

In our CLS assays, *cdt2Δ* and *rad24Δ* lived longer, while *lys9Δ* lifespan was the same as the wild type. This is opposite to Pombase annotation, which lists *cdt2Δ, rad24Δ* as short-lived and *lys9Δ* as long-lived. Those data come mainly from pooled, competitive screens and flow-cytometry–based viability measurements; pooled designs can be affected by outgrowth effects, whereas our non-competitive CFU assay avoids this issue and may explain the difference [67, 68]. Mechanistically, there is no evidence in the literature that *cdt2* is involved in TORC1 regulation. In addition, *rad24* promotes Wee1 stability in fission yeast, and TORC1 inhibition reduces Wee1 levels; thus, a *rad24* deletion could further destabilize Wee1, but a fall in Wee1 has not been shown to inhibit TORC1 in *S. pombe* [69]. Thus, we cannot yet link the longer CLS of *cdt2Δ/rad24Δ* to TORC1 inhibition [38]. Finally, although *cdt2Δ* and *rad24Δ* combine H_2_O_2_ resistance with longer CLS in our study, *lys9Δ* does not, indicating that stress resistance and longevity often coincide, but not always.

Altogether, we have identified a large number of mutants exhibiting resistance to hydrogen peroxide using our dynamic screen. In addition, we found 15 mutants that grow well (or even better) in oxidative stress conditions, thus, decoupling growth from stress gene expression programmes. Although the exact mechanisms behind this resistance are not within the scope of this manuscript, this study enhances our knowledge base of resistant mutants. Expanding our efforts beyond hydrogen peroxide, we are now able to engineer fission yeast to resist different types of stress and utilise them towards biotechnological applications [42, 44]. Our results show that following growth over time can help uncover meaningful phenotypes that may be missed in single-time-point screens. While a single late timepoint recovers a majority of resistant mutants (457), time-resolved analysis was essential to identify a substantial fraction (∼30%) of resistance phenotypes (a total of 661). The results also suggest that oxidative stress resistance is not always linked to growth rates and CLS and that different pathways may support survival in distinct ways. This work provides a practical method for identifying oxidative stress-related genes and offers a clear starting point for deeper investigation into the cellular mechanisms that help fission yeast manage environmental stress.

## Materials and Methods

### Media preparation

Yeast was grown in YES medium (3% glucose) throughout. To prevent contamination during the 4-day-long imaging course G418 was added. Three conditions were used throughout: control, 10 mM H_2_O_2_, and 12 mM H_2_O_2_. Hydrogen peroxide (30% w/v; Sigma-Aldrich) was added to molten medium to achieve the desired concentrations.

### Library preparation and phenotypic screening

50 mL media have been used for each Singer RoToR plus plate. A copy of the *S. pombe* Bioneer deletion library (version 5; 3,420 mutants) was arrayed in 384-format on YES plates using a RoToR robot (Singer Instruments) and incubated for 48 h at 32 °C. These served as templates for pinning onto control and H_2_O_2_-containing plates. Pinning was performed at 3% pressure with short 384-pins and no source mixing to minimize biomass, as dense colonies show increased stress tolerance [56]. Plates were incubated at 32 °C for and phenotypes were monitored for 4 days. Colony growth was imaged at 0, 20, 24, 44, 48, 68, 72, 92, and 96 h using an Epson Perfection V850 Pro scanner (400 dpi, reflective mode). Images were pre-processed for analysis with the R package Gitter [46], quantifying colony areas. Quantification data were linked to fission yeast gene IDs based on the plate geography.

### Data analysis

Colony size data were normalized to 0 h (growth fold-change) and to the corresponding control (resistance ratio). Data visualization and statistical tests were performed in RStudio using *ggplot2* [70]. Resistance ratios between 10 mM and 12 mM H_2_O_2_ were compared using the Wilcoxon rank-sum test. Heatmaps of growth patterns were generated in Morpheus [47], and Gene Ontology (GO) enrichment of fast-growing mutants was assessed using Metascape [50]. Fifteen mutants exhibited resistance ratios >1.15 under both peroxide concentrations at all time points. Using *k*-means clustering (Morpheus), mutants were grouped into seven clusters, and average growth fold-change across time was plotted. Mutants that grew slowly in control but rapidly under 12 mM H_2_O_2_ were prioritized for validation.

### Validation of resistance to hydrogen peroxide by spot assay

Candidate mutants and wild-type strains were grown in YES broth to mid-log phase (OD_600_ = 0.5), incubated for 30 min, and serially diluted 10-fold. Dilutions were spotted on YES plates containing 0–3 mM H_2_O_2_ and incubated at 32 °C for 48 h. Growth was imaged using a Vilber Lourmat FUSION Solo S system. *Pyp1Δ* [51] and *gsk3Δ* [28] strains served as positive controls.

### Chronological Lifespan Assay

Mutants were cultured in YES broth at 32 °C until stationary phase. Daily serial dilutions were plated on YES agar to determine colony-forming units (CFUs). Day 0 CFUs were defined as 100% survival, and subsequent counts were expressed relative to this value. CLS curves were generated from three biological replicates, with at least three technical replicates per time point.

## Acknowledgments

This work was supported by funding to C.R. from the Biotechnology and Biological Sciences Research Council [Research grant numbers: BB/V006916/1, BB/V006916/2]. C.R. also acknowledges support and funding of the group from the Medical Research Council [Grant number: MR/W001462/1].

## Supplemental Tables

**Supplemental Table 1**. List of all strains and their resistance ratios across all timepoints examined in 10 mM hydrogen peroxide.

**Supplemental Table 2**. List of all strains and their resistance ratios across all timepoints examined in 12 mM hydrogen peroxide.

**Supplemental Table 3**. List of the 661 strains with a resistance ratio above 1.1 in one or more of the timepoints examined.

## References

1. Chen D, Wilkinson CRM, Watt S, Penkett CJ, Toone WM, Jones N, and Bähler J (2008). Multiple pathways differentially regulate global oxidative stress responses in fission yeast. Mol Biol Cell. 19(1): 308–317. doi: 10.1091/MBC.E07-08-0735,.

2. López-Maury L, Marguerat S, and Bähler J (2008). Tuning gene expression to changing environments: from rapid responses to evolutionary adaptation. Nat Rev Genet. 9(8): 583–593. doi: 10.1038/NRG2398.

3. Kovacic P, and Somanathan R (2013). Broad overview of oxidative stress and its complications in human health. Open J Prev Med. 3(1): 32–41. doi: 10.4236/ojpm.2013.31005.

4. Finkel T, and Holbrook NJ (2000). Oxidants, oxidative stress and the biology of ageing. Nature. 408(6809): 239–247. doi: 10.1038/35041687;KWRD=SCIENCE.

5. Rhee SG (2006). H2O2, a necessary evil for cell signaling. Science (1979). 312(5782): 1882–1883. doi: 10.1126/SCIENCE.1130481/ASSET/DF68F3B6-E317-4CB0-8CA1-1D3A1ECF39AD/ASSETS/GRAPHIC/1882-1.GIF.

6. Veal EA, Day AM, and Morgan BA (2007). Hydrogen Peroxide Sensing and Signaling. Mol Cell. 26(1): 1– 14. doi: 10.1016/J.MOLCEL.2007.03.016/ASSET/843F2CCC-DC85-4640-A57E-E1A62A2E9FC3/MAIN.ASSETS/GR5.JPG.

7. Hoffman CS, Wood V, and Fantes PA (2015). An ancient yeast for young geneticists: A primer on the Schizosaccharomyces pombe model system. Genetics. 201(2): 403–423. doi: 10.1534/GENETICS.115.181503/-/DC1.

8. Wood V et al. (2002). The genome sequence of Schizosaccharomyces pombe. Nature. 415(6874): 871– 880. doi: 10.1038/NATURE724,.

9. Kim DU et al. (2010). Analysis of a genome-wide set of gene deletions in the fission yeast Schizosaccharomyces pombe. Nat Biotechnol. 28(6): 617–623. doi: 10.1038/NBT.1628,.

10. Roux PP, and Blenis J (2004). ERK and p38 MAPK-Activated Protein Kinases: a Family of Protein Kinases with Diverse Biological Functions. Microbiology and Molecular Biology Reviews. 68(2): 320– 344. doi: 10.1128/MMBR.68.2.320-344.2004,.

11. Bahn YS, Xue C, Idnurm A, Rutherford JC, Heitman J, and Cardenas ME (2007). Sensing the environment: Lessons from fungi. Nat Rev Microbiol. 5(1): 57–69. doi: 10.1038/NRMICRO1578,.

12. González A, and Hall MN (2017). Nutrient sensing and TOR signaling in yeast and mammals. EMBO J. 36(4): 397–408. doi: 10.15252/EMBJ.201696010,.

13. Laplante M, and Sabatini DM (2012). MTOR signaling in growth control and disease. Cell. 149(2): 274–293. doi: 10.1016/j.cell.2012.03.017.

14. Valvezan AJ, and Manning BD (2019). Molecular logic of mTORC1 signalling as a metabolic rheostat. Nat Metab. 1(3): 321–333. doi: 10.1038/S42255-019-0038-7,.

15. Rabanal-Ruiz Y, and Korolchuk VI (2018). mTORC1 and nutrient homeostasis: The central role of the lysosome. Int J Mol Sci. 19(3). doi: 10.3390/IJMS19030818,.

16. Jewell JL, Russell RC, and Guan KL (2013). Amino acid signalling upstream of mTOR. Nat Rev Mol Cell Biol. 14(3): 133–139. doi: 10.1038/NRM3522,.

17. Fernandes SA, Angelidaki DD, Nüchel J, Pan J, Gollwitzer P, Elkis Y, Artoni F, Wilhelm S, Kovacevic-Sarmiento M, and Demetriades C (2024). Spatial and functional separation of mTORC1 signalling in response to different amino acid sources. Nat Cell Biol. 26(11): 1918–1933. doi: 10.1038/S41556-024-01523-7,.

18. Chen D, Toone WM, Mata J, Lyne R, Burns G, Kivinen K, Brazma A, Jones N, and Bähler J (2003). Global Transcriptional Responses of Fission Yeast to Environmental Stress. Mol Biol Cell. 14(1): 214. doi: 10.1091/MBC.E02-08-0499.

19. Shiozaki K, and Russell P (1995). Counteractive roles of protein phosphatase 2C (PP2C) and a MAP kinase kinase homolog in the osmoregulation of fission yeast. EMBO J. 14(3): 492. doi: 10.1002/J.1460-2075.1995.TB07025.X.

20. Shiozaki K, and Russell P (1996). Conjugation, meiosis, and the osmotic stress response are regulated by Spc1 kinase through Atf1 transcription factor in fission yeast. Genes Dev. 10(18): 2276–2288. doi: 10.1101/GAD.10.18.2276,.

21. Wilkinson MG, Samuels M, Takeda T, Mark Toone W, Shieh JC, Toda T, Millar JBA, and Jones N (1996). The Atf1 transcription factor is a target for the Sty1 stress-activated MAP kinase pathway in fission yeast. Genes Dev. 10(18): 2289–2301. doi: 10.1101/GAD.10.18.2289,.

22. Toone WM, Kuge S, Samuels M, Morgan BA, Toda T, and Jones N (1998). Regulation of the fission yeast transcription factor Pap1 by oxidative stress: requirement for the nuclear export factor Crm1 (Exportin) and the stress-activated MAP kinase Sty1/Spc1. Genes Dev. 12(10): 1453. doi: 10.1101/GAD.12.10.1453.

23. Vivancos AP, Jara M, Zuin A, Sansó M, and Hidalgo E (2006). Oxidative stress in Schizosaccharomyces pombe: Different H 2O2 levels, different response pathways. Molecular Genetics and Genomics. 276(6): 495–502. doi: 10.1007/S00438-006-0175-Z,.

24. Bozonet SM, Findlay VJ, Day AM, Cameron J, Veal EA, and Morgan BA (2005). Oxidation of a eukaryotic 2-Cys peroxiredoxin is a molecular switch controlling the transcriptional response to increasing levels of hydrogen peroxide. Journal of Biological Chemistry. 280(24): 23319–23327. doi: 10.1074/jbc.M502757200.

25. Veal EA, Findlay VJ, Day AM, Bozonet SM, Evans JM, Quinn J, and Morgan BA (2004). A 2-Cys peroxiredoxin regulates peroxide-induced oxidation and activation of a stress-activated MAP kinase. Mol Cell. 15(1): 129–139. doi: 10.1016/j.molcel.2004.06.021.

26. Ohmiya R, Kato C, Yamada H, Aiba H, and Mizuno T (1999). A fission yeast gene (prr1+) that encodes a response regulator implicated in oxidative stress response. J Biochem. 125(6): 1061–1066. doi: 10.1093/OXFORDJOURNALS.JBCHEM.A022387,.

27. Kennedy PJ, Vashisht AA, Hoe KL, Kim DU, Park HO, Hayles J, and Russell P (2008). A Genome-Wide Screen of Genes Involved in Cadmium Tolerance in Schizosaccharomyces pombe. Toxicological Sciences. 106(1): 124. doi: 10.1093/TOXSCI/KFN153.

28. Rallis C, Townsend S, and Bähler J (2017). Genetic interactions and functional analyses of the fission yeast gsk3 and amk2 single and double mutants defective in TORC1-dependent processes. Sci Rep. 7. doi: 10.1038/SREP44257,.

29. Piotrowski JS, Simpkins SW, Li SC, Deshpande R, McIlwain SJ, Ong IM, Myers CL, Boone C, and Andersen RJ (2015). Chemical genomic profiling via barcode sequencing to predict compound mode of action. Methods Mol Biol. 1263: 299. doi: 10.1007/978-1-4939-2269-7_23.

30. Barazandeh M, Kian Gaikani H, Pattanshetti R, Uche Ogbede J, Sinha S, Carr CE, Giaever G, Nislow C, and Author C (2025). Bar-seq: A robust, platform-agnostic method for massively parallel cell-based screens. G3 Genes|Genomes|Genetics. doi: 10.1093/G3JOURNAL/JKAF166.

31. Rodríguez-López M, Bordin N, Lees J, Scholes H, Hassan S, Saintain Q, Kamrad S, Orengo C, and Bähler J (2023). Broad functional profiling of fission yeast proteins using phenomics and machine learning. Elife. 12. doi: 10.7554/ELIFE.88229.

32. Hibbing ME, Fuqua C, Parsek MR, and Peterson SB (2010). Bacterial competition: surviving and thriving in the microbial jungle. Nat Rev Microbiol. 8(1): 15. doi: 10.1038/NRMICRO2259.

33. Kamrad S, Rodríguez-Ló Pez M, Cotobal C, Correia-Melo C, Ralser M, and Rg BäHler J Pyphe, a python toolbox for assessing microbial growth and cell viability in high-throughput colony screens. doi: 10.7554/eLife.55160.

34. Pickering AM, Vojtovich L, Tower J, and Davies KJA (2012). Oxidative Stress Adaptation with Acute, Chronic and Repeated Stress. Free Radic Biol Med. 55: 109. doi: 10.1016/J.FREERADBIOMED.2012.11.001.

35. Chen BR, and Runge KW (2009). A new Schizosaccharomyces pombe chronological lifespan assay reveals that caloric restriction promotes e?cient cell cycle exit and extends longevity. Exp Gerontol. 44(8): 493–502. doi: 10.1016/j.exger.2009.04.004.

36. Lu C, Brauer MJ, and Botstein D (2009). Slow growth induces heat-shock resistance in normal and respiratory-deficient yeast. Mol Biol Cell. 20(3): 891–903. doi: 10.1091/MBC.E08-08-0852,.

37. Brauer MJ, Huttenhower C, Airoldi EM, Rosenstein R, Matese JC, Gresham D, Boer VM, Troyanskaya OG, and Botstein D (2008). Coordination of Growth Rate, Cell Cycle, Stress Response, and Metabolic Activity in Yeast. Mol Biol Cell. 19(1): 352. doi: 10.1091/MBC.E07-08-0779.

38. Rodríguez-López M, Gonzalez S, Hillson O, Tunnacliffe E, Codlin S, Tallada VA, Bähler J, and Rallis C (2020). The GATA Transcription Factor Gaf1 Represses tRNAs, Inhibits Growth, and Extends Chronological Lifespan Downstream of Fission Yeast TORC1. Cell Rep. 30(10): 3240-3249.e4. doi: 10.1016/j.celrep.2020.02.058.

39. Powers RW, Kaeberlein M, Caldwell SD, Kennedy BK, and Fields S (2006). Extension of chronological life span in yeast by decreased TOR pathway signaling. Genes Dev. 20(2): 174–184. doi: 10.1101/GAD.1381406.

40. Roux AE, Quissac A, Chartrand P, Ferbeyre G, and Rokeach LA (2006). Regulation of chronological aging in Schizosaccharomyces pombe by the protein kinases Pka1 and Sck2. Aging Cell. 5(4): 345–357. doi: 10.1111/J.1474-9726.2006.00225.X,.

41. MacLean M, Harris N, and Piper PW (2001). Chronological lifespan of stationary phase yeast cells; a model for investigating the factors that might influence the ageing of postmitotic tissues in higher organisms. Yeast. 18(6): 499–509. doi: 10.1002/YEA.701.

42. Nielsen J (2012). Production of biopharmaceutical proteins by yeast Advances through metabolic engineering. doi: 10.4161/bioe.22856.

43. Das PK, Sahoo A, and Veeranki VD (2024). Recombinant monoclonal antibody production in yeasts: Challenges and considerations. Int J Biol Macromol. 266(Pt 2). doi: 10.1016/j.ijbiomac.2024.131379.

44. Mohd Azhar SH, Abdulla R, Jambo SA, Marbawi H, Gansau JA, Mohd Faik AA, and Rodrigues KF (2017). Yeasts in sustainable bioethanol production: A review. Biochem Biophys Rep. 10: 52–61. doi: 10.1016/J.BBREP.2017.03.003.

45. Rallis C, and Bähler J (2016). Cell-based screens and phenomics with fission yeast. Crit Rev Biochem Mol Biol. 51(2): 86–95. doi: 10.3109/10409238.2015.1103205.

46. Wagih O, and Parts L (2014). gitter: A Robust and Accurate Method for Quantification of Colony Sizes From Plate Images. G3 Genes|Genomes|Genetics. 4(3): 547–552. doi: 10.1534/G3.113.009431.

47. Wood V, Harris MA, McDowall MD, Rutherford K, Vaughan BW, Staines DM, Aslett M, Lock A, Bähler J, Kersey PJ, and Oliver SG (2012). PomBase: a comprehensive online resource for fission yeast. Nucleic Acids Res. 40(D1): D695–D699. doi: 10.1093/NAR/GKR853.

48. Malecki M, Bitton DA, Rodríguez-López M, Rallis C, Calavia NG, Smith GC, and Bähler J (2016). Functional and regulatory profiling of energy metabolism in fission yeast. Genome Biology 2016 17:1. 17(1): 1–18. doi: 10.1186/S13059-016-1101-2.

49. Starruß J, De Back W, Brusch L, and Deutsch A (2014). Morpheus: a user-friendly modeling environment for multiscale and multicellular systems biology. Bioinformatics. 30(9): 1331–1332. doi: 10.1093/BIOINFORMATICS/BTT772.

50. Zhou Y, Zhou B, Pache L, Chang M, Khodabakhshi AH, Tanaseichuk O, Benner C, and Chanda SK (2019). Metascape provides a biologist-oriented resource for the analysis of systems-level datasets. Nat Commun. 10(1): 1–10. doi: 10.1038/S41467-019-09234-6;SUBJMETA=114,1314,2164,2391,2401,631;KWRD=CELLULAR+SIGNALLING+NETWORKS,DATA+INTEGRATION,DATA+MINING,DATA+PROCESSING.

51. Boronat S, Marte L, Vega M, García-Santamarina S, Cabrera M, Ayté J, and Hidalgo E (2020). The Hsp40 Mas5 Connects Protein Quality Control and the General Stress Response through the Thermo-sensitive Pyp1. iScience. 23(11). doi: 10.1016/j.isci.2020.101725.

52. Maldonado E, Morales-Pison S, Urbina F, and Solari A (2023). Aging Hallmarks and the Role of Oxidative Stress. Antioxidants 2023, Vol 12, Page 651. 12(3): 651. doi: 10.3390/ANTIOX12030651.

53. Rallis C, López-Maury L, Georgescu T, Pancaldi V, and Bähler J (2014). Systematic screen for mutants resistant to TORC1 inhibition in fission yeast reveals genes involved in cellular ageing and growth. Biol Open. 3(2): 161–171. doi: 10.1242/BIO.20147245.

54. Chen D, Wilkinson CRM, Watt S, Penkett CJ, Toone WM, Jones N, and Bähler J (2008). Multiple Pathways Differentially Regulate Global Oxidative Stress Responses in Fission Yeast. Mol Biol Cell. 19(1): 308. doi: 10.1091/MBC.E07-08-0735.

55. Váchová L, and Palková Z (2018). How structured yeast multicellular communities live, age and die? FEMS Yeast Res. 18(4): 33. doi: 10.1093/FEMSYR/FOY033.

56. Stewart PS, Roe F, Rayner J, Elkins JG, Lewandowski Z, Ochsner UA, and Hassett DJ (2000). Effect of Catalase on Hydrogen Peroxide Penetration into Pseudomonas aeruginosa Biofilms. Appl Environ Microbiol. 66(2): 836. doi: 10.1128/AEM.66.2.836-838.2000.

57. Ralser M, Wamelink MM, Kowald A, Gerisch B, Heeren G, Struys EA, Klipp E, Jakobs C, Breitenbach M, Lehrach H, and Krobitsch S (2007). Dynamic rerouting of the carbohydrate flux is key to counteracting oxidative stress. J Biol. 6(4). doi: 10.1186/JBIOL61,.

58. Shenton D, Smirnova JB, Selley JN, Carroll K, Hubbard SJ, Pavitt GD, Ashe MP, and Grant CM (2006). Global translational responses to oxidative stress impact upon multiple levels of protein synthesis. Journal of Biological Chemistry. 281(39): 29011–29021. doi: 10.1074/jbc.M601545200.

59. Nakashima A, Sato T, and Tamanoi F (2010). Fission yeast TORC1 regulates phosphorylation of ribosomal S6 proteins in response to nutrients and its activity is inhibited by rapamycin. J Cell Sci. 123(5): 777. doi: 10.1242/JCS.060319.

60. Wilkinson MG, Samuels M, Takeda T, Mark Toone W, Shieh JC, Toda T, Millar JBA, and Jones N (1996). The Atf1 transcription factor is a target for the Sty1 stress-activated MAP kinase pathway in fission yeast. Genes Dev. 10(18): 2289–2301. doi: 10.1101/GAD.10.18.2289.

61. Petersen J, and Russell P (2016). Growth and the Environment of Schizosaccharomyces pombe. Cold Spring Harb Protoc. 2016(3): pdb.top079764. doi: 10.1101/PDB.TOP079764.

62. Olin-Sandoval V, Yu JSL, Miller-Fleming L, Alam MT, Kamrad S, Correia-Melo C, Haas R, Segal J, Peña Navarro DA, Herrera-Dominguez L, Méndez-Lucio O, Vowinckel J, Mülleder M, and Ralser M (2019). Lysine harvesting is an antioxidant strategy and triggers underground polyamine metabolism. Nature 2019 572:7768. 572(7768): 249–253. doi: 10.1038/s41586-019-1442-6.

63. Lopez-Girona A, Furnari B, Mondesert O, and Russell P (1999). Nuclear localization of Cdc25 is regulated by DNA damage and a 14-3-3 protein. Nature. 397(6715): 172–175. doi: 10.1038/16488,.

64. López-Avilés S, Grande M, González M, Helgesen AL, Alemany V, Sanchez-Piris M, Bachs O, Millar JBA, and Aligue R (2005). Inactivation of the Cdc25 phosphatase by the stress-activated Srk1 kinase in fission yeast. Mol Cell. 17(1): 49–59. doi: 10.1016/J.MOLCEL.2004.11.043/ATTACHMENT/5CD09326-B69B-4EE3-A137-6C98ECBD40B7/MMC4.DOC.

65. Havens CG, and Walter JC (2011). Mechanism of CRL4Cdt2, a PCNA-dependent E3 ubiquitin ligase. Genes Dev. 25(15): 1568. doi: 10.1101/GAD.2068611.

66. Zhang JM, Zheng JX, Ding YH, Zhang XR, Suo F, Ren JY, Dong MQ, and Du LL (2020). CRL4Cdt2 ubiquitin ligase regulates Dna2 and Rad16 (XPF) nucleases by targeting Pxd1 for degradation. PLoS Genet. 16(7): e1008933. doi: 10.1371/JOURNAL.PGEN.1008933.

67. Zahedi Y, Durand-dubief M, and Ekwall K (2020). High-Throughput Flow Cytometry Combined with Genetic Analysis Brings New Insights into the Understanding of Chromatin Regulation of Cellular Quiescence. International Journal of Molecular Sciences 2020, Vol 21, Page 9022. 21(23): 9022. doi: 10.3390/IJMS21239022.

68. Vega M, Castillo D, de Cubas L, Wang Y, Huang Y, Hidalgo E, and Cabrera M (2022). Antagonistic effects of mitochondrial matrix and intermembrane space proteases on yeast aging. BMC Biol. 20(1): 1– 18. doi: 10.1186/S12915-022-01352-W/FIGURES/7.

69. Alao JP, Johansson-Sjölander J, Rallis C, and Sunnerhagen P (2020). Caffeine Stabilises Fission Yeast Wee1 in a Rad24-Dependent Manner but Attenuates Its Expression in Response to DNA Damage. Microorganisms. 8(10): 1512. doi: 10.3390/MICROORGANISMS8101512.

70. Wickham H (2016). ggplot2. doi: 10.1007/978-3-319-24277-4.

